# Slow but flexible or fast but rigid? Discrete and continuous processes compared

**DOI:** 10.1101/2023.08.20.554008

**Authors:** Matteo Priorelli, Ivilin Peev Stoianov

## Abstract

A tradeoff exists when dealing with complex tasks composed of multiple steps. High-level cognitive processes can find the best sequence of actions to achieve a goal in uncertain environments, but they are slow and require significant computational demand. In contrast, lower-level processing allows reacting to environmental stimuli rapidly, but with limited capacity to determine optimal actions or to replan when expectations are not met. Through reiteration of the same task, biological organisms find the optimal tradeoff: from action primitives, composite trajectories gradually emerge by creating task-specific neural structures. The two frameworks of active inference – a recent brain paradigm that views action and perception as subject to the same *free energy minimization* imperative – well capture high-level and low-level processes of human behavior, but how task specialization occurs in these terms is still unclear. In this study, we compare two strategies on a dynamic pick-and-place task: a hybrid (discrete-continuous) model with planning capabilities and a continuous-only model with fixed transitions. Both models rely on a hierarchical (intrinsic and extrinsic) structure, well suited for defining reaching and grasping movements, respectively. Our results show that continuous-only models perform better and with minimal resource expenditure but at the cost of less flexibility. Finally, we propose how discrete actions might lead to continuous attractors and compare the two frameworks with different motor learning phases, laying the foundations for further studies on bio-inspired task adaptation.

## 1 Introduction

How does the brain support the efficient execution of tasks comprising multiple steps, such as picking and placing an object? While a sequence of movements is easy to plan in static contexts, a difficulty emerges when acting in dynamic environments, e.g., when the object has to be grasped on the fly. Tackling complex multi-step tasks is known to occur deep in the cortical hierarchy, by areas processing discrete, slow-varying entities [1]. But the advantage of discrete representations useful for planning [2] comes at a cost: if an object moves too fast, deep processing might be too slow for the grasping action to succeed.

Motor skill consolidation is a well-known phenomenon [3], clearly evident in athletes [4, 5, 6]: during repeated exposure to a task, an initial learning phase gives way to autonomous movements as the athlete becomes more proficient [7]. Cortical involvement is gradually reduced as specialized neural structures are constructed in lower, fast subcortical regions [8], including fine adjustment in the spinal cord reflexes [9]. These structures cannot extract invariant representations to perform high-level decision-making but they react rapidly to the sensorium. The basal ganglia are known to be central to task specialization as the striatum encodes chunked representations of action steps composing a learned habit [10, 11]. But this change seems much more pervasive, involving information processing within the cortex itself and a general activity shift from anterior to posterior regions [12], as evidenced by plastic changes in the primary motor cortex [13]. The brain mechanisms underlying such changes are still unclear, and understanding their computational basis is compelling.

An interesting proposal is that the brain maintains a model of the task dynamics, and skill consolidation consists of fine-tuning this model to achieve a stable behavior accounting for environmental uncertainty [14, 15]. This hypothesis fits with a recent theory called *active inference*, which brings insights of increasing appeal into the computational role of the nervous system [16, 17, 18]. Active inference provides a formalization of the two essential components of control, sensing and acting, which supposedly aim to resolve the critical goal of all organisms: to survive in uncertain environments by operating within preferred states (e.g., maintaining a constant temperature). As in predictive coding [19], active inference assumes that organisms perceive the environment through an internal generative model constructed by inferring how hidden causes produce sensations [17, 20]. Crucially, perception and action jointly minimize a quantity called *free energy*: while perception gradually adjusts internal expectations to match sensory evidence, action gradually samples those sensations that make the expectations true [17, 20].

The two frameworks of this theory have been used to analyze high-level and low-level processes of human behavior under a unified perspective. Active inference in continuous time defines an internal dynamic model about the self and external targets in generalized coordinates of motion (position, velocity, acceleration, and so on) [20, 21]. Elementary movements are then resolved by local suppression of proprioceptive prediction errors in classical reflex arcs [22], forcing a physical change in the actuators. This framework is ideal for interacting with real environments; however, it falls short when it comes to multi-step actions, since free energy minimization only deals with current sensory signals.

At the other extreme, decision-making is addressed by the discrete formulation [23, 24, 25], which views planning as an inferential mechanism [26, 27]. Discrete active inference models leverage Partially Observed Markov Decision Processes (POMDPs) to plan abstract actions over (yet unobserved) outcomes thanks to the minimization of the *expected free energy*, i.e., the free energy that the agent expects to perceive in the future [28, 29, 30, 31]. This framework seems well suited to analyze cortical operations [1], with higher biological plausibility compared to reinforcement learning [32, 33]. However, the downside of discrete computations is the lack of real-time interaction with the environment.

Between the two stands a third, *hybrid*, framework, which specifies a discrete model combined with its continuous counterpart [34, 35]. This involves the transformation of continuous signals into discrete messages through Bayesian Model Reduction [36, 37]. Solving multi-step tasks with a hybrid model implies associating elementary continuous trajectories to discrete states for planning. The mathematical link between prediction errors in continuous formulations and variational free energy in discrete formulations is straightforward: precision-weighted prediction errors are variational free energy gradients, which are minimized at free energy minima, where the gradients are destroyed. This approach has not received adequate attention yet, as not many implementations can be found in the literature [2, 18, 34, 38, 35, 39, 40], none simulating applications in dynamic contexts.

Importantly, it is not entirely clear how active inference could simulate the virtuous cycle of motor skill acquisition, which begins with the rigid information processing of subcortical structures and culminates with the emergence of task-specific structures. To shed light on this issue, the hybrid framework could be highly relevant: discrete computations can simulate the behavior of an agent that has no prior idea of what actions to take. However, as soon as the agent has learned a policy that corresponds to a sequence of a priori-defined actions, it can adapt by encoding the transitions between discrete states into continuous dynamics, which then represent a learned motor skill.

In summary, this paper foregrounds the distinction between active inference under continuous and hybrid generative models, with special reference to their neurobiological implementation and the biomimetic advantages afforded by both kinds of models. Several perspectives might help understand the key distinction. Active inference under continuous state-space models can be regarded as a generalization of “control as inference”; e.g., [41, 42, 43]. Control as inference itself can be thought of as generalizing linear quadratic control to include nonlinear and deep (hierarchical) state space models. In neurobiology, this kind of control is often referred to in terms of equilibrium point or trajectory hypotheses [44, 45]; namely, using motor reflexes to eliminate proprioceptive or actuator prediction errors – where the predictions provide setpoints or trajectories that replace motor commands. In the active inference literature, these schemes are sometimes referred to as “merely reflexive” and are apt to describe various forms of homeostatic behavior or behaviors driven by autonomous dynamics (e.g., central pattern generators). Merely reflexive active inference should be compared with the “planning as inference” [46, 26, 35] afforded by equipping continuous state space models with discrete state-spaces that include the consequences of sequential policies. This brings a future-pointing aspect to behavior, afforded by rolling out sequences of actions and evaluating them under posterior predictive distributions over outcomes. In neurobiology, this can be framed as a move from homeostasis to allostasis [47, 39] and affords the capacity for information-seeking and preference-seeking behaviors that go beyond merely reflexive behavior. The price paid for this prospective kind of active inference is the computational cost and requisite course graining (into discrete states). This trade-off speaks to the integration of continuous and discrete models of the sort described below.

## 2 Methods

### 2.1 Perception, control, and the variational free energy

Active inference assumes that organisms perceive the environment through an internal generative model that infers how external causes produce sensations and how they evolve [17, 20]. In the continuous domain, this model is usually factorized into probability distributions over hidden states 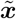, hidden causes 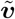 (i.e., causal variables over the hidden states dynamics), and observable outcomes 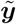:

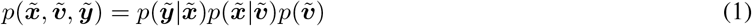

The symbol ∼ indicates variables encoded in generalized coordinates of motion, representing instantaneous trajectories (position, velocity, acceleration, and so on), e.g., 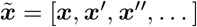. These distributions are assumed to be Gaussian:

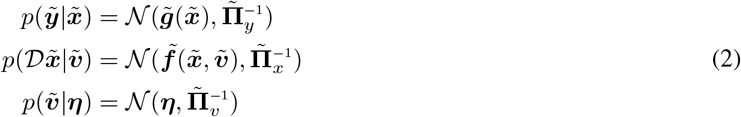

corresponding to the following non-linear stochastic equations that represent how the environment evolves:

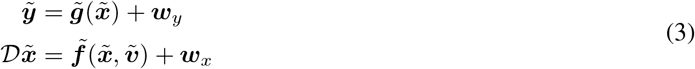

Here, 𝒟 is the differential shift operator that shifts all the temporal orders by one, i.e.: 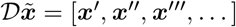, and we indicated with the symbol 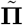 the generalized precisions (or inverse variances) of the distributions. Directly evaluating the posterior 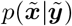 is intractable since it involves the computation of the inaccessible marginal 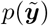. The variational solution is to approximate the posteriors with *recognition distributions* of a more tractable form. Perceptual inference then turns into a minimization of the difference between the approximate and real posteriors, which can be formalized in terms of a KL divergence – equivalent to the expectation, over the approximate posterior, of the difference between the two log-probabilities:

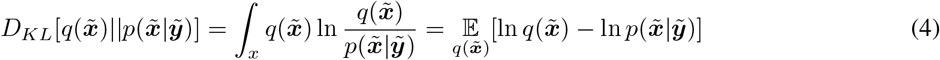

Given that the denominator 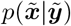 still depends on the marginal 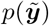, we express the KL divergence in terms of the log-evidence and the *Variational Free Energy* (VFE), and minimize the latter quantity instead:

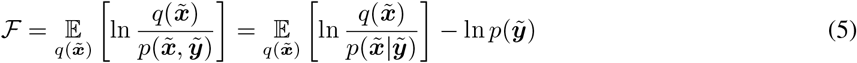

Since the KL divergence is non-negative (due to Jensen’s inequality), the VFE provides an upper bound on surprise – thus, its optimization improves both evidence and model fit. One common assumption about the form of the recognition distributions is the Laplace approximation [16]:

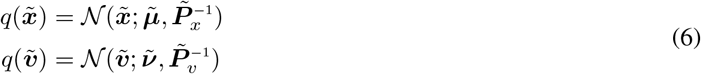

where the parameters 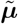 and 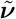 are called *beliefs* over hidden states and hidden causes, while 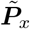 and 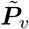 are their precisions. Under this assumption, the free energy breaks down to a simple formula (see [18] for more details):

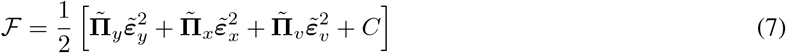

where 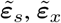 and 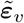 are respectively prediction errors of sensations, dynamics and prior:

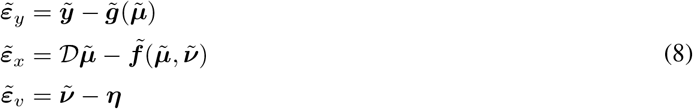

and we included in *C* the terms that disappear when computing the gradients. Then, minimizing the free energy with respect to hidden states and hidden causes turns into an iterative parameter update through the following expressions:

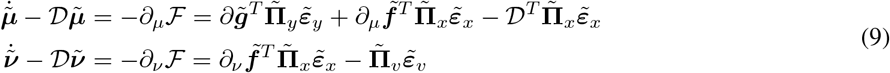

Notably, the VFE can also be minimized by acting, computing the following motor control signals:

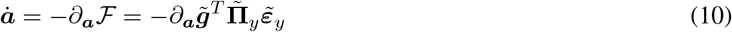

In continuous-time active inference, goals are generally encoded as prior beliefs in the hidden causes 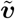. These generate sensory predictions about a specific evolution of the world in accordance with the agent’s beliefs. If these predictions are realized by movement, the agent eventually looks for those states that make his or her belief true. This process – known as *self-evidencing* – allows implementing goal-directed behavior through VFE minimization, keeping the agent in predictable and safer spaces [17]. Note that this process can be scaled up to construct a hierarchical model, wherein each level is involved in computing a prediction, comparing it with the state below, and updating its state with the generated prediction error. These simple steps are repeated throughout the whole hierarchy so that, from a prior belief encoding a goal, a cascade of proprioceptive trajectories is generated that is eventually realized by the lowest levels of the motor system.

### 2.2 Planning with the expected free energy

Although active inference in continuous time can deal with real-world problems by keeping track of instantaneous trajectories, it has several limitations and a narrow use, as it cannot easily handle more general types of actions arising from decision-making. VFE minimization can only adjust the internal generative model depending on current (or past) observations, and it does not evaluate future states and outcomes. To endow an agent with this ability, a quantity called *Expected Free Energy* (EFE) is considered [23, 24]. We consider a generative model similar to Equation 1, but controlled by policies ***π***:

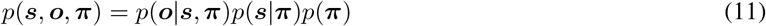

where ***s*** and ***o*** are discrete hidden states and outcomes. Note that policies are not simple stimulus-response mappings – as in Reinforcement Learning schemes – but sequences of actions. The agent’s generative model is factorized as in POMDPs:

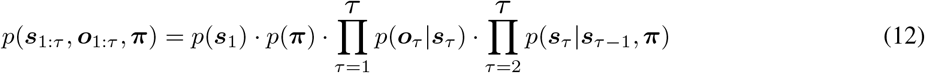

where 𝒯 is the total number of discrete steps. These elements are represented with categorical distributions:

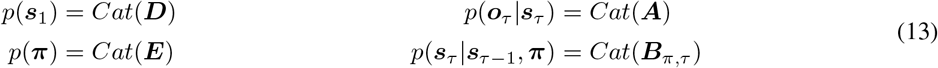

where ***D*** encodes beliefs about the initial state, ***E*** encodes the prior over policies, ***A*** is the likelihood matrix and ***B***_*π, τ*_ is the transition matrix. Similar to the continuous case, perceptual inference relies on an approximate posterior distribution *q*(***s***_1: *τ*_, ***π***), and minimizes the VFE – which in turn minimizes the Kullback-Leibler (KL) divergence between the approximate and real posteriors:

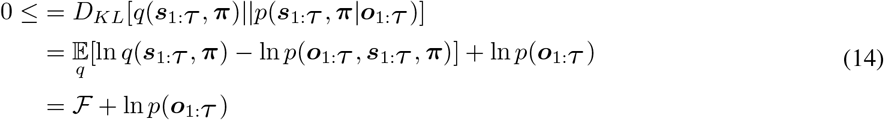

If we assume, under the mean-field approximation, that the approximate posterior factorizes into independent distributions:

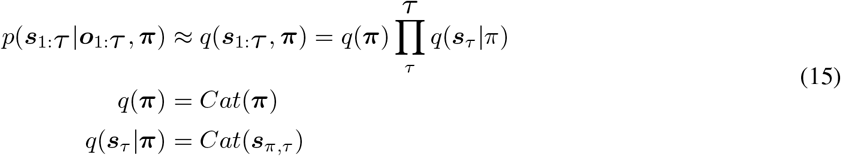

we can use the standard technique of variational message passing to infer each posterior *q*(***s***_*τ*_ |***π***) and combine them into a global posterior *q*(***s***_1: *τ*_ |***π***). In order to update the posterior about hidden states, we combine the messages from past states, future states, and outcomes, express each term with its sufficient statistics, and finally apply a softmax function to get a proper probability distribution:

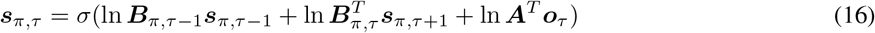

Instead, action planning involves inferring those policies that lead to the desired outcomes. Hence, the EFE is specifically constructed by considering future outcomes as random variables, and by conditioning over them:

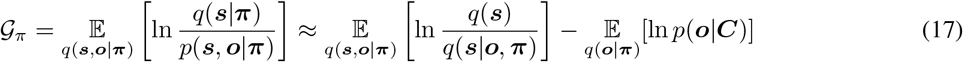

where the probability distribution *p*(***o***|***C***) encodes preferred outcomes. The last two RHS terms are respectively called *epistemic* (uncertainty-reducing) and *pragmatic* (goal-seeking). In order to update *q*(***π***), we combine the messages from the prior over policies given by the matrix ***E***, and from future observations conditioned upon policies; we can approximate the latter by the EFE conditioned at a particular time *τ* :

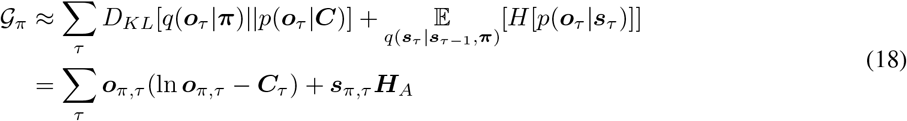

where:

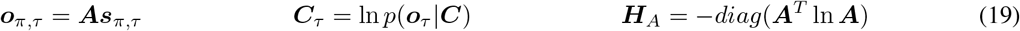

Finally, we select the action *u* that is the most likely under all policies:

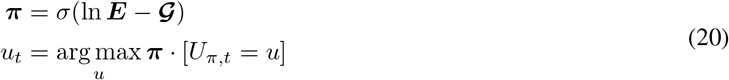

As continuous-time active inference, discrete models can be scaled as well. This leads to a hierarchy of temporal dynamics, wherein low levels adapt more frequently to sensory observations, while high levels construct increasingly invariant representations of the environment. Differently from continuous models, active inference in discrete state-spaces allows to operate with abstract actions at different levels, creating a hierarchical plan composed of several subgoals [2].

### 2.3 Bayesian model reduction in hybrid models

To get a discrete model working with the richness of continuous signals, a form of communication with the continuous framework is required. Since the two frameworks operate in different domains, we need a way to obtain a continuous prior from a discrete prediction and, concurrently, to estimate a discrete outcome based on continuous evidence. Both problems can be easily addressed through Bayesian model reduction [37, 36]. Consider a generative model *p*(***θ, y***) with parameters ***θ*** and data ***y***:

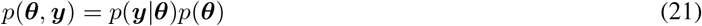

and an additional distribution 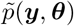, which is a reduced version of the first model if the likelihood of some data is the same under both models and the only difference rests upon the specification of the priors 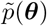. We can express the posterior of the reduced model as:

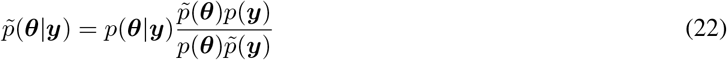

The procedure for computing the reduced posterior is the following: first, we integrate over the parameters to obtain the evidence ratio of the two models:

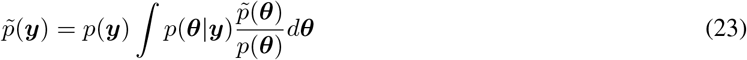

Then, we define an approximate posterior *q*(***θ***) and we compute the reduced VFE in terms of the full model:

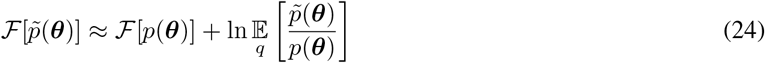

Correspondingly, the approximate posterior of the reduced model can be written in terms of the full model:

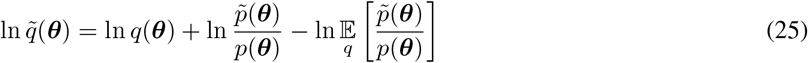

In order to apply this technique to the communication between discrete and continuous models, we first consider the hidden causes ***v*** as parameters and denote with ***y*** the continuous observations. We then assume that the full prior probability depends on a discrete outcome:

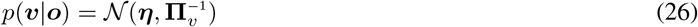

In this way, the parameter ***η*** acts as a prior over the hidden causes. We also assume that the agent maintains *M* (Gaussian) reduced models:

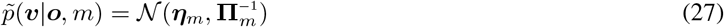

with prior ***η***_*m*_. These priors correspond to the agent’s hypotheses about the causes of its sensorium, i.e., they encode a specific configuration of the continuous model, and each of them is associated with a particular discrete outcome *o*_*τ,m*_, i.e., ***o***_*τ*_ = [*o*_*τ*,1_, …, *o*_*τ,M*_]. The latter are computed by marginalizing ***o***_*π,τ*_ over all policies:

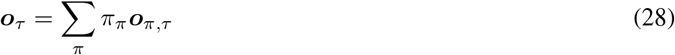

Then, the prior ***η*** of the full model, which represents the actual agent’s parameters of the current state of the world, is obtained by simply performing a Bayesian model average between every specified model:

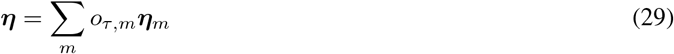

thus transforming discrete probabilities into a continuous value. As a result, this quantity biases the belief ***ν*** over hidden causes following Equations 9-8. Similarly, having defined the full and reduced approximate posteriors:

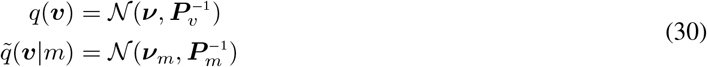

we compute an ascending message (acting as an observation for the discrete model) through Bayesian model comparison, i.e., by comparing the agent’s expectations with the observed evidence. More formally, the free energy computed in Equation 24 acts as a hint to how well a reduced model explains continuous observations, and it corresponds to the prior surprise plus the log-evidence for each outcome model accumulated over time:

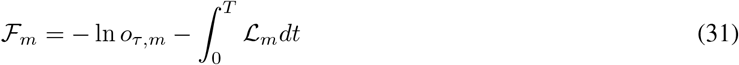

We refer to *T* as sampling time window: every discrete time step *τ* corresponds to a continuous period *T* during which evidence is accumulated. The Laplace approximation [48] leads to a simple form of the second RHS term:

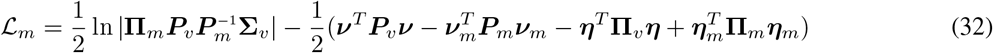

where the mean and precision of the *m*th reduced model are:

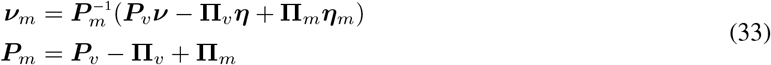

After having computed the free energy associated with each model, they are normalized through a softmax function to get a proper probability, which is finally used to estimate the current discrete state:

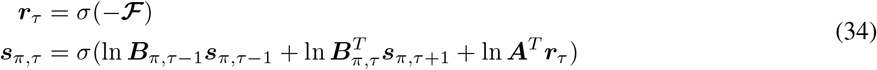

In summary, each value of ***r***_*τ*_ indicates how probable an agent’s hypothesis is based on the accumulated continuous evidence. In contrast to the discrete-only case, the discrete actions in a hybrid model are only implicitly used to compute the posterior over policies, and the link between the two frameworks is possible by associating each discrete outcome model *o*_*τ,m*_ to a specific continuous prior ***η***_*m*_. These priors are combined to realize an average of the agent’s hypotheses, and are concurrently compared with the estimated continuous trajectories to infer the true cause of the sensorium.

## 3 Results

To shed light on task specialization under active inference, we first delineate how to specify multiple hidden state dynamics in the agent’s generative model. These dynamics are related to the agent’s desires (e.g., reaching or grasping an object) or *intentions* [49, 50]. The defined method is used as the basis for tackling dynamic multi-step tasks in both hybrid or continuous-only frameworks, showing that discrete actions and continuous trajectories work under similar mechanisms. We then compare the behavioral differences between hybrid and continuous frameworks in terms of planning capabilities and reaction times. To this aim, we consider a pick-and-place operation. The agent’s body is an 8-DoF simulated arm with two fingers. At each trial, a target object spawns with a random and unknown position; the agent’s goal is to reach the object, grasp it, reach a random goal position, and open the fingers to place the object. To assess the model’s capabilities in dynamic environments, at each trial a velocity with random direction is assigned to the object, so that the grasping action might fail if the object moves too fast.

### 3.1 Flexible intentions for dynamic multi-step tasks

Whether an agent maintains a simple continuous generative model or makes use of a discrete model for planning, to operate in realistic scenarios it needs to specify beliefs over environmental entities. Maintaining such additional beliefs has been used to actively infer object-centric representations [51, 52] or for simulating oculomotor behavior [53]; here, we turn to the problem of tackling dynamic tasks that comprise multiple steps or composite actions. If the agent’s goal is to reach a moving object, it must maintain an estimate of its hand position as well as of the object position [54]. Designing a dynamics function with an attractor toward a static prior configuration is not helpful in dynamic contexts, and directly embedding exteroceptive observations of the object (which are low-level representations) in the internal model is not biologically plausible, since it would require to define the sensory mapping in the model dynamics as well.

Formally, we consider *beliefs* ***µ*** about different environmental entities operating in the same domain, i.e., ***µ*** = [***µ***_1_, …, ***µ***_*N*_], where *N* is the number of entities. When we talk about beliefs, we mean the expected value of hidden or latent states generating observations. We assume that these beliefs generate observations ***y*** in parallel, through a likelihood function ***g***(***µ***) with the same factorization:

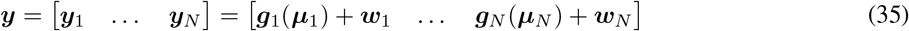

where the letter *w* indicates a (Gaussian) noise term. Each belief can then be inferred by inverting this mapping:

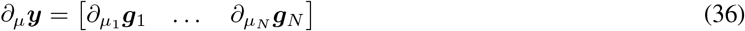

and by following the update rules defined in Equation 9. In this way, the agent can track the trajectory of its body along with the trajectories of moving objects or other agents. At this point, the dynamic nature of a task can be tackled through the definition of appropriate *intentions*, or functions that generate a possible future state by combining the beliefs over some environmental entities [54]:

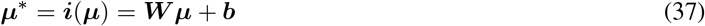

Here, we specified a form that could be realized through a simple combination of neurons (although more advanced behaviors can be realized with nonlinear functions). In particular, the matrix ***W*** performs a linear transformation of the beliefs, affording dynamic behavior since every entity is constantly updated through the observations. On the other hand, the vector ***b*** imposes a static configuration, e.g., a fixed position to reach. Note that differently from the likelihood function ***g***(***µ***), the function ***i***(***µ***) manipulates and combines all the environmental entities.

To clarify how Equation 37 works, it is useful to model the pick-and-place operation considered before. To focus on the control aspects, we assume a simulated agent endowed with the following simplified sensory modalities: (i) a proprioceptive observation of the arm’s joint angles (with dimension *n*_*p*_ = 8); (ii) a visual observation encoding the positions of hand, object, and goal (with dimension *n*_*v*_ = 3); (iii) a tactile observation:

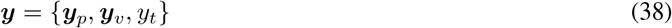

For simplicity, we directly provide the 2D Cartesian coordinates of the hand, object, and goal as visual input, i.e., ***y***_*v*_ = [***y***_*v,a*_, ***y***_*v,o*_, ***y***_*v,g*_]. Tactile observations are defined through a Boolean function and inform the agent whether all the fingers – the last four joints – touch an external object. Here, we only consider the Cartesian position of the last level (i.e., the hand). However, a more realistic scenario should comprise information about all intermediate limb positions, which may be used to infer the correct posture from exteroceptive sensations only [55].

Although limb trajectories are generated by *intrinsic* kinematic information (e.g. joint angles), the movements that the agent wants to realize are usually expressed in an *extrinsic* domain (e.g., 3D visual space). These two continuous modalities are needed in pick-and-place operations because reaching movements are better described in an extrinsic reference frame, while opening and closing the hand are operations better understood from an intrinsic reference frame. We thus consider beliefs in both intrinsic (***µ***_*i*_) and extrinsic (***µ***_*e*_) domains. It is also convenient to express in these two domains the objects the agent interacts with. In this way, for each object the agent can automatically infer a possible joint configuration, which can be further restricted by appropriate priors (e.g., a particular grip depending on the object’s affordances). We further assume that the agent maintains a belief over limb lengths ***µ***_*l*_ in order to compute the extrinsic Cartesian coordinates of the hand and every object. Here, this belief is kept fixed, although it is possible to infer it [55, 56]. Finally, we also keep a belief over tactile observations *µ*_*t*_. Summing up:

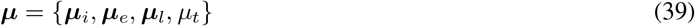

where the first two beliefs comprise components about the arm, object, and goal, reflecting the decomposition of the visual sensory signal:

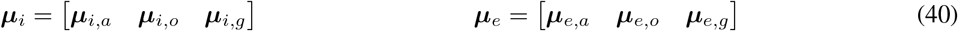

Since joint angles generate Cartesian coordinates with a one-to-one mapping, it is natural to place the intrinsic belief at the top of this hierarchy, and define the following likelihood function ***g***_*e*_:

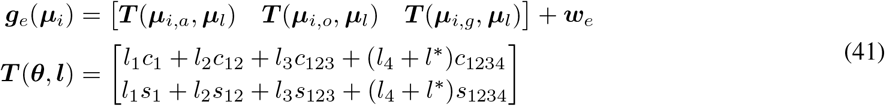

where ***T*** (***θ, l***) is the forward kinematics. As a result, the intrinsic belief is found by inferring the most likely kinematic configuration that may have generated the extrinsic belief [55]; this naturally performs inverse kinematics without explicitly requiring it in the dynamics function of the intrinsic belief. Here, *l*_*n*_ is the length of the *n*th segment, and we used a compact notation to indicate the sine and cosine of the sum of angles. Since we are not interested in the positions of the fingers during the reaching task, the kinematic likelihood function only computes the Cartesian position of the hand, which is found by extending the length of the last limb by a grasping distance *l*^*\**^. Finally, proprioceptive predictions are generated through a mapping ***g***_*p*_ that extracts the arm’s joint angles from the intrinsic belief, while exteroceptive sensations – position and touch – are generated by an identity function:

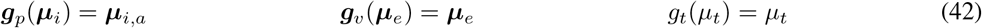

The belief dynamics can account for many factors including friction, gravity, etc. – but for simplicity we define them as a combination of the agent’s intentions. For a pick-and-place task, it is useful to specify an intention for each step: (i) reach the object; (ii) reach the goal position; (iii) close the fingers to grasp the object; (iv) open the fingers to release the object. The first two intentions (indexed by a second subscript) can be realized in an extrinsic reference, setting the hand position equal to the inferred object/goal positions:

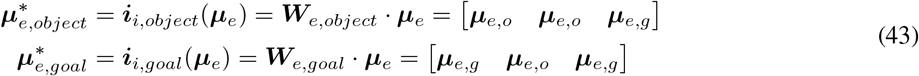

with the help of the following transformations:

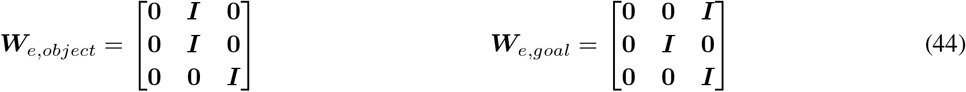

where the elements **0** and ***I*** are 2×2 zero and identity matrices, corresponding to the x and y coordinates. As explained later, the extrinsic belief over the hand, steered toward one of the environmental entities, generates a prediction error that propagates back to the intrinsic level, inferring a kinematic configuration that is realized by acting. Intentions with a similar form can also be defined at the intrinsic level, setting the arm’s joint angles equal to the inferred object/goal configurations (recall that the agent constantly maintains potential configurations related to the entities): this results in faster dynamics as the intentions directly operate on the intrinsic domain required for movement.

Grasping/placing intentions are accomplished by generating a future intrinsic belief with the fingers in a fixed closed/open configuration:

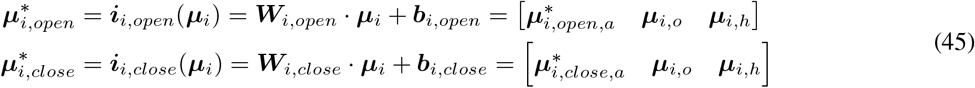

Here, ***W***_*i,open*_ and ***W***_*i,close*_ ensure that the first 4 components of the arm (thus, excluding the fingers) maintain the current configuration:

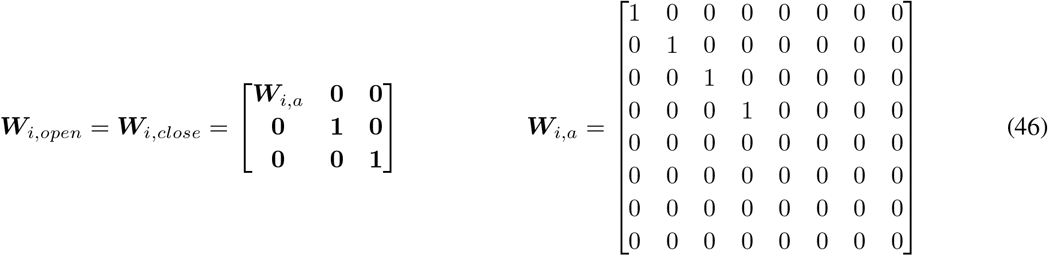

where the elements **0** and ***I*** are 8×8 zero and identity matrices, corresponding to the arm’s joint angles. Instead, ***b***_*i,open*_ and ***b***_*i,close*_ impose closed/open angles *θ*_*c*_ and *θ*_*o*_ to the fingers:

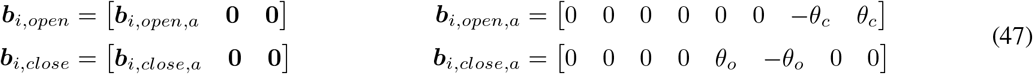

where **0** is a zero vector. In summary, we defined two sets of competing intentions operating in different domains: reaching an object or reaching a goal position (in extrinsic reference frames), and closing or opening the hand (in intrinsic reference frames). Figure 1 provides a graphical representation of the link between environmental beliefs and flexible intentions. In the following, we turn to the definition of the two (hybrid and continuous-only) control methods, showing how these intentions are used in practice. The difference between the two methods regards where intentions are embedded and how they affect the system dynamics.

**Figure 1:**
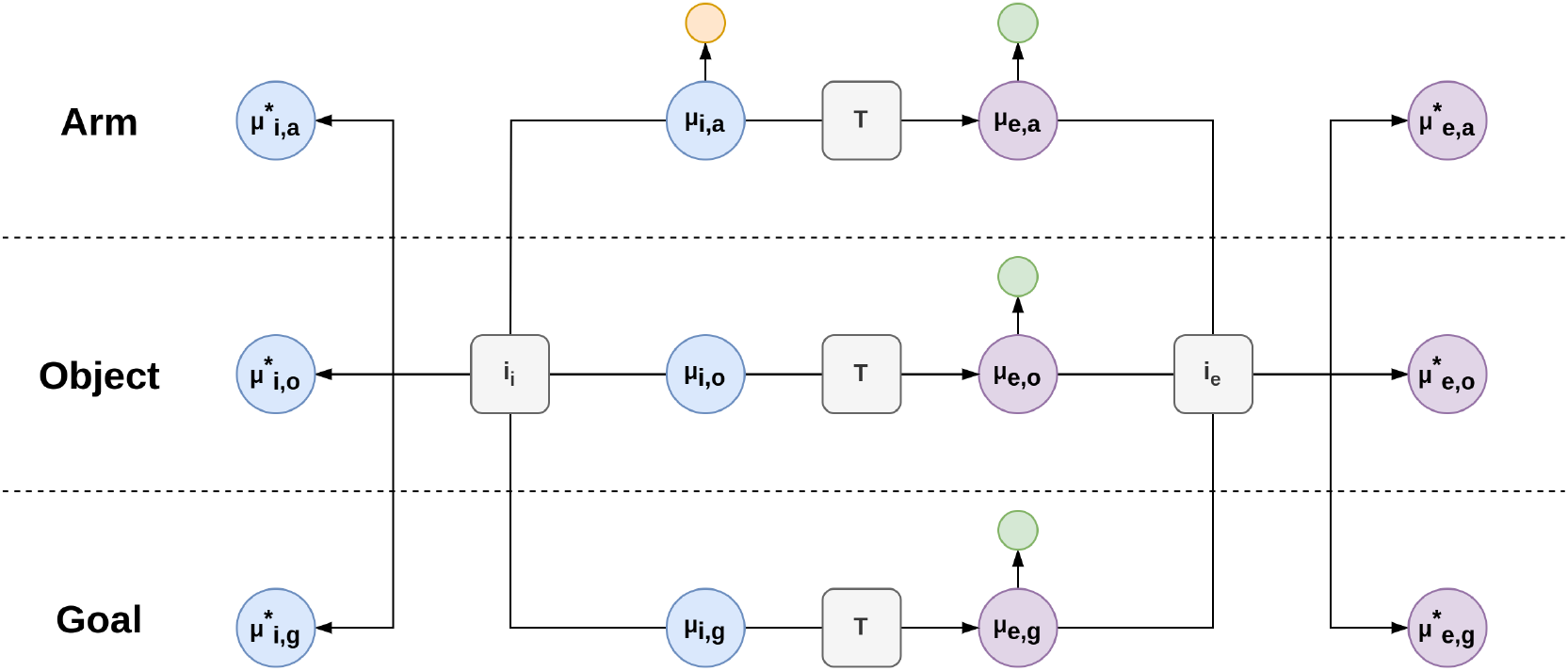
Graphical representation of the relationship between environmental beliefs and intentions. Intrinsic and extrinsic beliefs are linked by a function ***T*** computing forward kinematics. Both beliefs have components representing the arm, an object, and a goal position. Orange and green circles represent proprioceptive and visual observations, respectively (note that only the arm component generates proprioceptive predictions). Future states are computed through intentions ***i***_*i*_ and ***i***_*e*_ that manipulate and combine all the environmental entities from a given domain.

### 3.2 Hybrid models and discrete goals

In this section, we delineate how the defined intentions can be used to simulate a pick-and-place operation with a hybrid model, which we then compare to a specific phase of motor skill learning. Designing a discrete model involves the specification of discrete states ***s***, a likelihood matrix ***A***, a transition matrix ***B***, and actions ***u***. In the task considered, the discrete hidden states encode: (i) whether the hand and object are in the same position, in different positions, or both in the goal position; and (ii) whether the hand is open, closed or has grasped the object. In total, these factors combine in 9 possible process states. There are two likelihood matrices: (i) a matrix ***A*** performs a simple identity mapping from the discrete to the continuous model; and (ii) a matrix ***A***_*t*_ returns a discrete tactile observation ***o***_*t*_ – encoded by a Bernoulli distribution – signaling whether or not the object is grasped (hence, the continuous observation *y*_*t*_ is not used). The transition matrix ***B*** is defined such that the object can be grasped only when both hand and object are in the same position. Note that every state has a certain probability – e.g., depending on sampling time window *T* – that the transition might fail. Finally, the discrete actions ***u*** are equivalent to the intentions defined before, with the addition of a *stay* action. The trial is successful only if the agent has placed the object in the goal position and the hand is open.

For the discrete and continuous models to communicate, each discrete outcome has to be associated with a reduced continuous prior. Since the latter is usually static, to deal with a dynamic environment we let it depend on a combination of the intentions. This has the advantage that if the reduced priors are generated at each discrete time step, a correct configuration over the hidden states can be imposed even with a moving object – whose position will be dynamically inferred by the continuous model. However, note that a richer design would be to link the reduced priors to the hidden causes and encode the intentions in the latter [57]. With this method, dynamic inference of discrete variables emerges naturally by generating the reduced priors from different dynamics functions maintained by the agent. Crucially, this means that not just the position, but the whole instantaneous trajectory is used for inference and planning.

Since the agent maintains intrinsic and extrinsic beliefs, a single discrete outcome *o*_*τ,m*_ generates two different sets of reduced priors, ***η***_*i,m*_ and ***η***_*e,m*_. For example, if the hand is open and in the object position, the reduced priors are:

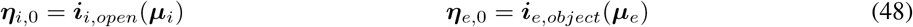

Note that some outcomes generate the same reduced priors (e.g., the open-hand condition does not affect the extrinsic prior), or do not impose any bias over the hidden states (e.g., if the hand and the object are in different positions, the mapping of the extrinsic level reduces to an identity and the corresponding reduced prior will be equal to the posterior). Through Bayesian Model Average (BMA), the full priors ***η***_*i*_ and ***η***_*e*_ are then computed, which act over the continuous models as prior prediction errors ***ε***_*η,i*_ and ***ε***_*η,e*_:

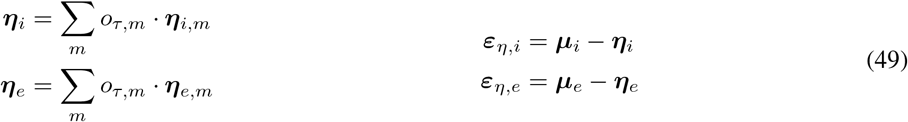

Finally, the updates for the hidden states are found by the following update rules:

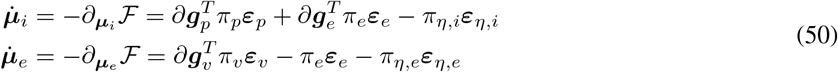

Where ℱ is the variational free energy of the continuous models, ***ε***_*p*_, ***ε***_*e*_, ***ε***_*v*_ are respectively the proprioceptive, extrinsic, and visual prediction errors, and we indicated with the letter *π* the precisions of the related likelihood functions. Note that both updates have a similar form, with the only differences lying in the type of sensory observations, and in the direction of the extrinsic prediction error (backward vs forward). In particular, the gradient ∂***g***_*e*_ performs kinematic inversion from intrinsic to extrinsic beliefs [55].

On the other side, ascending messages are found by comparing, for a *sampling time window T*, the models of both intrinsic and extrinsic posteriors through the free energies 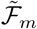 of the reduced models:

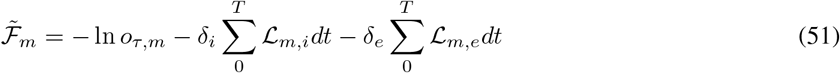

where ℒ_*m,i*_ and ℒ_*m,e*_ are the log-evidences of intrinsic and extrinsic beliefs – computed via Equation 32 – respectively weighted by gains *δ*_*i*_ and *δ*_*e*_. The latter quantities modulate the contributions of different domains. Here, they are kept fixed throughout the tasks but it is assumed that higher values result in faster reaction times and evidence accumulation during perceptual decision-making [58]. Note that a long enough sampling time window *T* is needed to accumulate evidence from the lower modalities and plan the next movement, as will be clear later. At this point, the discrete hidden states are inferred by combining the likelihoods of both the ascending message ***r***_*τ*_ and the tactile observation ***o***_*t*_:

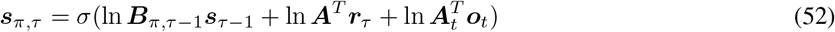

similar to Equation 34. In summary, the discrete model first computes policy probabilities depending on some prior preferences, and combines them to obtain a discrete outcome. The latter is used to compute – through flexible intentions – the reduced priors, which are further weighted to get a final configuration that the continuous models have to realize. This configuration acts as a prior over the 0th-order (i.e., position) hidden states, and the belief update breaks down to the computation of sensory gradients and prediction errors. Note that the extrinsic belief ***µ***_*e*_ can be biased through two different pathways: directly from the reduced prior of the discrete model, or indirectly through forward kinematics of the intrinsic belief ***µ***_*i*_. Correspondingly, the latter can be changed either by imposing a particular configuration in the intrinsic domain through the discrete model, or by the backward pathway of the extrinsic belief, through computation of the gradient ∂***g***_*e*_ [55].

What are the implications from a biological perspective? It has been hypothesized that motor skill learning is roughly composed of three different phases [59]: a first cognitive phase where movements are inefficient and one has to consciously explore different strategies to achieve a goal; an associative stage where the learner becomes more proficient and transitions between movements are more fluid; a final autonomous phase where the learner has reached almost complete autonomy from conscious task-specific thinking and can move without cognitive effort. Although the reality is likely to be far more complex than this simple decomposition, with different degrees of proficiency corresponding to specific neural processing, it may be useful for analyzing the key principles that underlie the change in neural activity about task specialization. In this view, active inference provides interesting insights about this process because: (i) it builds upon an internal generative model of the task dynamics, which can be constantly refined by experience; (ii) it assumes that information processing starts and unfolds through the same principle of prediction error minimization, either with continuous or discrete representations. Under this perspective, the hybrid model described can be compared with the first cognitive phase – shown in Figure 2 – wherein the agent has not yet developed an appropriate automatism for pick-and-place operations, and has to constantly plan the next action to take based on the new evidence observed. The resulting goal-directed behavior is composed of simple action primitives (reaching, picking, or placing), which is inefficient and requires non-negligible energy demands.

**Figure 2:**
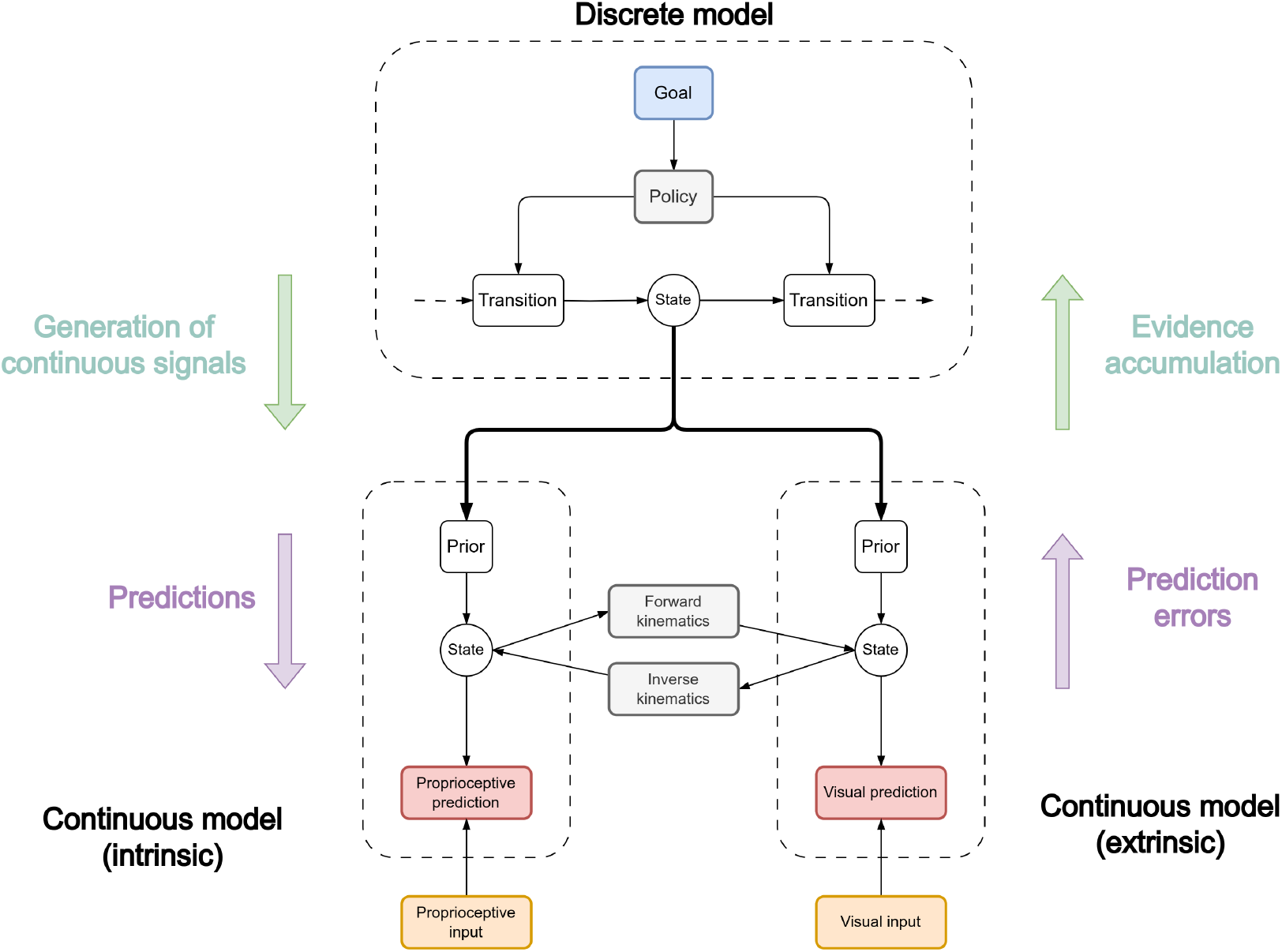
Graphical representation of the first cognitive phase. In the first trials of a novel task, the state transitions generally encode elementary actions, such as reaching, grasping, and releasing. The cognitive effort needed by highlevel processes to learn the correct sequence of elementary actions translates to the task dynamics being only visible in the discrete model, while the continuous levels only receive a prior representation (e.g., target states) and their job is mostly to realize the corresponding trajectories. This is in line with the hypothesis that in the first phase of motor learning a high cortical activity is recorded, especially in anterior (e.g., prefrontal) regions.

### 3.3 Continuous models and flexible intentions

Whether or not a discrete model is necessary depends on the nature of the task, even when considering actions composed of different steps. In some cases, an a-priori-defined sequence of movements (i.e., a habitual behavior) is all that is needed to solve a particular task, e.g., in rhythmic movements or, as in this case, a simple pick-and-place operation. Such scenarios may present little uncertainty about the order of elementary actions that does not necessarily involve repeated online decision-making.

But how can transitions between continuous trajectories be encoded in practice? To elucidate, we reveal a parallelism between high-level and low-level processes, shown in Figure 3a. In the discrete model, hidden states are conditioned on a policy that the agent can follow in a specific instant, and the overall state ***s***_*τ*_ is found by weighting the expected states from all policies. The same technique is applied to hybrid models, where the probability of every possible discrete outcome is combined with a continuous state to compute the average signal that will bias the lower levels. Along the same line, we propose that composite continuous dynamics can be generated by weighting the results from independent contributions, each encoding a simple trajectory. Specifically, we let the agent’s intentions depend on *hidden causes* ***v***, i.e., with beliefs ***ν***_*i*_ = [*ν*_*i*,1_, …, *ν*_*i,J*_] and ***ν***_*e*_ = [*ν*_*e*,1_, …, *ν*_*e,K*_], where *J* and *K* are the number of intrinsic and extrinsic intentions. Next, we factorize the prior distribution over the generalized beliefs 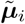 and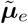 (in this case, up to the 1st order) into different probability distributions for each hidden cause:

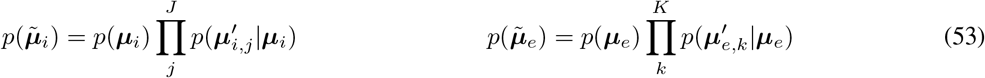

where:

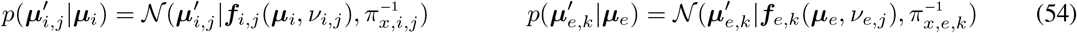

and *π*_*x,i,j*_ and *π*_*x,e,k*_ are the corresponding precisions. Each function acts as an attractive force proportional to the error between the current belief and desired states, the latter computed with the intentions defined before:

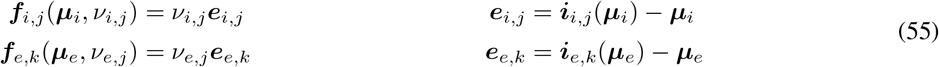

**Figure 3:**
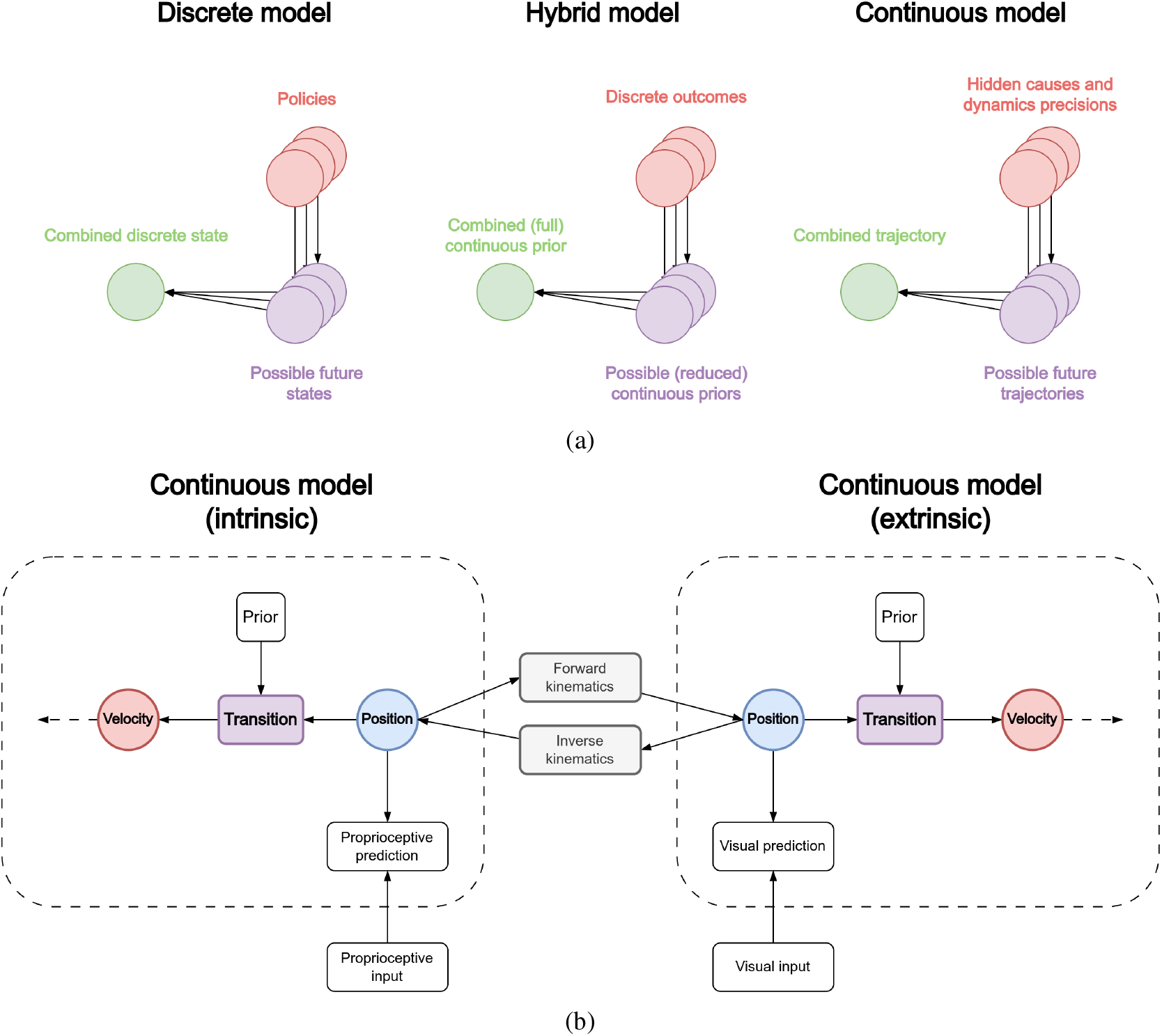
(a) Bayesian model average in discrete, hybrid, and continuous models. In discrete models, future states ***s***_*π,τ*_ are weighted by the probability of each policy *π*_*π*_, generating a combined discrete state ***s***_*τ*_. In hybrid models, continuous (reduced) priors ***η***_*m*_ are weighted by the probability of each discrete outcome modality *o*_*τ,m*_, generating a combined (full) continuous prior ***η***, which acts as a bias for the continuous model. In continuous models, future dynamic trajectories 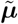 are weighted by the hidden causes ***v*** and dynamics precisions *π*_*x*_, generating a combined trajectory realized through movement. (b) Graphical representation of the third autonomous phase. The process of task specialization is complete. The discrete model – not shown here – has learned a representation abstract enough to include all the four main movements of the pick-and-place operation in a combined trajectory, and its role is only to command its execution and accumulate evidence from both modalities to infer when it has been completed.

Note that the hidden causes can either act as attractor gains, or specify the relative strength of one dynamics over the other [54, 60]; as a result, modulation of the hidden causes achieves a multi-step behavior. In fact, since the belief dynamics already store and embed every future goal, the belief follows the contribution of all active dynamics. In particular, we want the following behavior: (i) until the hand has not reached the object, execute only ***i***_*e,object*_ (i.e., reach the object) and ***i***_*i,open*_ (i.e., open the hand); (ii) as soon as the hand reaches the object, ***i***_*i,close*_ should start (i.e., close the hand); (iii) when the agent has grasped the object, the object-reaching intention should be replaced by the ***i***_*e,goal*_ (i.e., move the hand toward the goal position along with the object); (iv) when the agent has reached the goal position, execute ***i***_*i,open*_ again to release the object. Note that the object-reaching intention should be active until the agent has correctly grasped the object, since the latter may be moving and the grasping intention fail. The desired behavior can be easily implemented by computing and combining Boolean functions (or sigmoid functions to have smooth transitions) of the beliefs. Having defined the dynamics prediction errors for each intention:

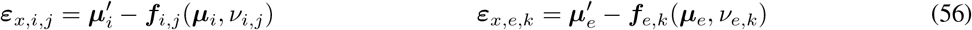

the update rules for the intrinsic and extrinsic beliefs will be a precision-weighted combination of sensory prediction errors and multiple attractor dynamics:

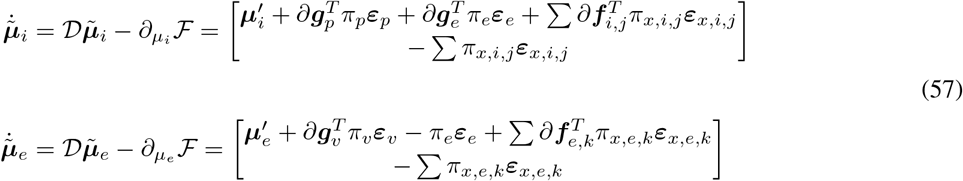

Note here the dynamics prediction errors acting as a forward message to the 1st-order of the beliefs. Finally, we update the beliefs over tactile sensations:

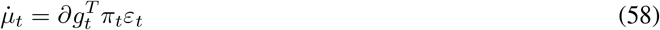

Figure 3b shows a graphical representation of the continuous-only model, compared to the third phase of motor learning. The continuous models now encode the dynamics of the entire task in the most efficient way because the mechanism is fully autonomous and does not require the repeated activation of higher levels for planning the next action. Differently than the previous model (Figure 2), which had to encode every possible elementary movement – either intrinsic or extrinsic – the increased efficiency is also the consequence of each modality encoding its dynamics independently from the others, translating to grasping (or reaching) movements being mapped mostly into the intrinsic (or extrinsic) modality. In fact, the dimensions of hidden causes *J* and *K* generally differ.

A sequence of time frames is shown in Figure 4a for a sample grasping trial with a continuous-only control. Note that the intrinsic and extrinsic beliefs comprise estimates of the arm, the object to be grasped, and the goal position. Defining continuous intentions in the intrinsic domain has the advantage that high hierarchical levels can impose priors not only over hand positions, but also over specific configurations of the arm and fingers. This is useful when the agent has to grasp an object in a particular manner depending on the task considered (e.g., power grip vs precision grip). Figure 4b shows a simple demonstration of this behavior, wherein the agent is required to grasp the object with a counterclockwise wrist rotation, and then place it in the goal position with a clockwise rotation. Here, there is a tradeoff between reaching the object and maintaining the imposed constraint on the wrist; this is evident in the middle of the movement when the agent waits until the object can be grasped with the correct rotation even if the latter is already in its peripersonal space.

**Figure 4:**
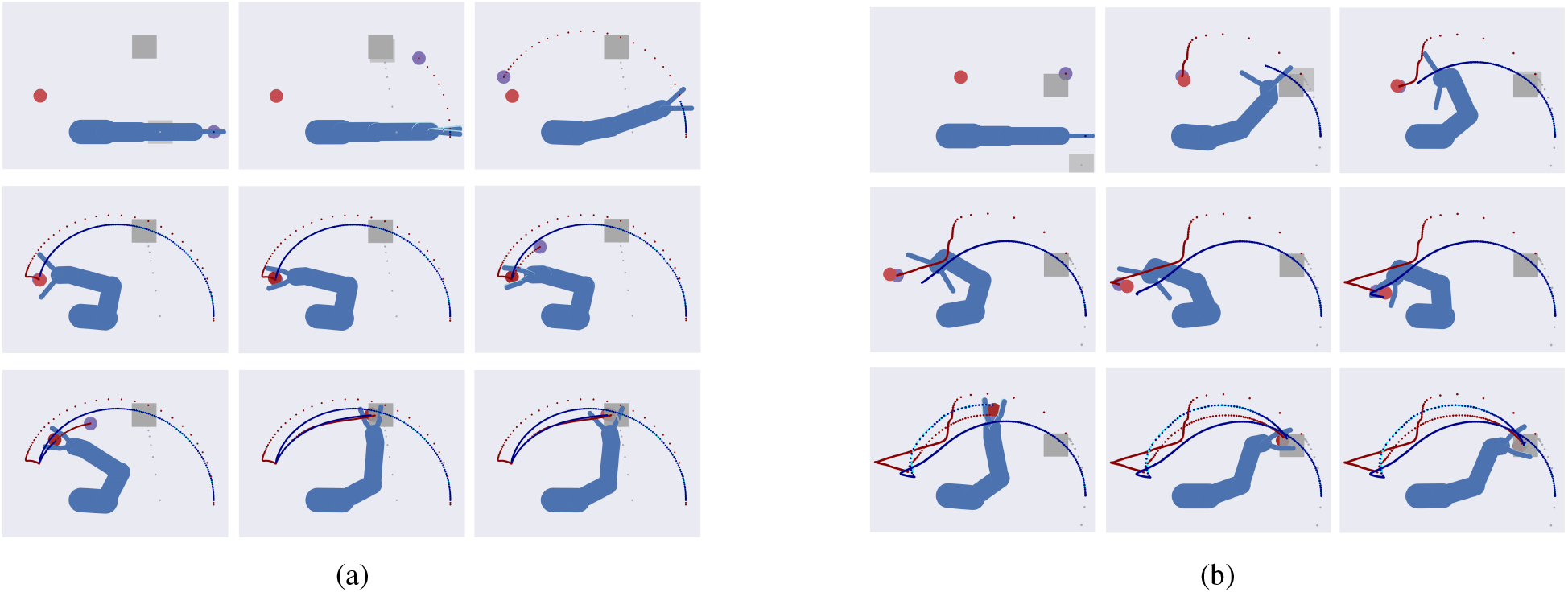
(a) Sequence of time frames of the grasping task with static object and goal. Real and estimated object positions are represented by red and purple circles, real and estimated goal positions by grey and dark grey squares, and real and estimated limb positions in blue and cyan. Hand, object, and goal belief trajectories are represented respectively with a blue, red, and grey dotted line. The object position is rapidly inferred while the hand reaches it. As soon as the object is grasped, the conditions over the hidden causes force the agent to switch dynamics; as a result, the object belief is pulled toward the goal position while the agent continues to track it. When the object is in the goal position, another transition between dynamics forces the agent to place the object. (b) Sequence of time frames of the grasping task with affordances and a moving object. The arm starts with the fingers closed and the object belief is initialized to the hand position. It starts tracking the object while slowly opening the hand and preparing a counterclockwise wrist rotation. Being subject to multi-domain constraints, this configuration is maintained until the moving object is grasped. When this happens, the agent rotates the wrist by 180 degrees clockwise and correctly places the object in the goal position.

### 3.4 Discrete and continuous processes compared

In summary, goal-directed behavior can be achieved through specular mechanisms operating at different hierarchical levels, as shown in Figure 3a. Comparing the inference of continuous and discrete hidden states, we note a relationship between the policy-dependent states ***s***_*π,τ*_ computed through the discrete transition matrix ***B***, and the trajectories 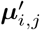 and 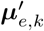 computed through the continuous intentions. While a policy-independent discrete state ***s***_*τ*_ is found via Bayesian model average of each policy probability *π*_*π*_, in the continuous case the hidden states ***µ***_*i*_ and ***µ***_*e*_ are computed by weighting each trajectory with the corresponding hidden causes *ν*_*i,j*_ and *ν*_*e,k*_ at each step of the task.

If we consider the hybrid model, we also note a similarity between the probability distributions of the reduced priors ***η***_*m*_ of Equation 27 and the probabilities of Equation 54 that compose the dynamics function. In the former case, each reduced prior is averaged through each outcome model probability *o*_*τ,m*_. In this case, however, the full prior biases the continuous belief through the overall prediction error *ε*_*η,i*_ (or *ε*_*η,e*_), which already contains the final configuration that the continuous model must realize until the next discrete step. In the continuous-only model, the intentions are instead used to compute different directions of update that independently pull the belief over the continuous hidden states. Also, the prior precision *π*_*η,i*_ of Equation 50 and the dynamics precisions *π*_*x,i,j*_ and *π*_*x,e,k*_ of Equation 57 play the same role of encoding the strength of the intention dynamics – that steers the belief toward a desired state – in contrast to the sensory likelihood – that keeps the agent close to what it is currently perceiving. Crucially, while the (intrinsic) weighted prediction error of the hybrid model is:

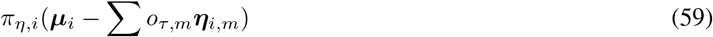

if the dynamics precisions have the same value *π*_*x,i*_, the combination of the weighted dynamics prediction error of the continuous-only model takes the form:

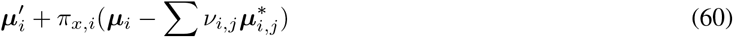

Hence, encoding different precisions for each intention permits an additional modulation. As a result, in the continuous-only model the future trajectories are subject to two different processes [57]. A *fast* process that imposes (and infers) future states based on the hidden causes ***ν***_*i*_ and ***ν***_*e*_; but also a *slow* process that learns the precisions *π*_*x,i,j*_ and *π*_*x,e,k*_ of every trajectory. This result lends itself to an intuitive interpretation: in active inference, a low precision in a specific sensory modality implies that it cannot be trusted and the agent should update its internal state relying on other signals. Correspondingly, a trajectory with high precision is a good option for minimizing prediction errors. In other words, the agent is more confident in using an intention either for solving the task considered or for explaining the current context – given the reciprocal interactions between action and perception.

In order to evaluate the capacity of the continuous and hybrid models to solve a grasping task in static and dynamic conditions, we first run an experiment with 9 target velocities ranging from 0 to 80 pixels per time step, 500 grasping trials per condition (i.e., 4500 trials in total per model). The sampling time window of the hybrid model – the period *T* in Equation 51 – was fixed at 50 time steps. The positions of the target and the goal, as well as the direction of the target, were randomly sampled at the beginning of every trial. Also, the target dimension was randomly chosen at each trial, within a defined range. For each condition, accuracy was computed as the average number of successful trials in which the agent successfully picked the object and placed it in the goal position before the deadline of 6000 time steps. The simulation results are displayed in Figure 5a.

**Figure 5:**
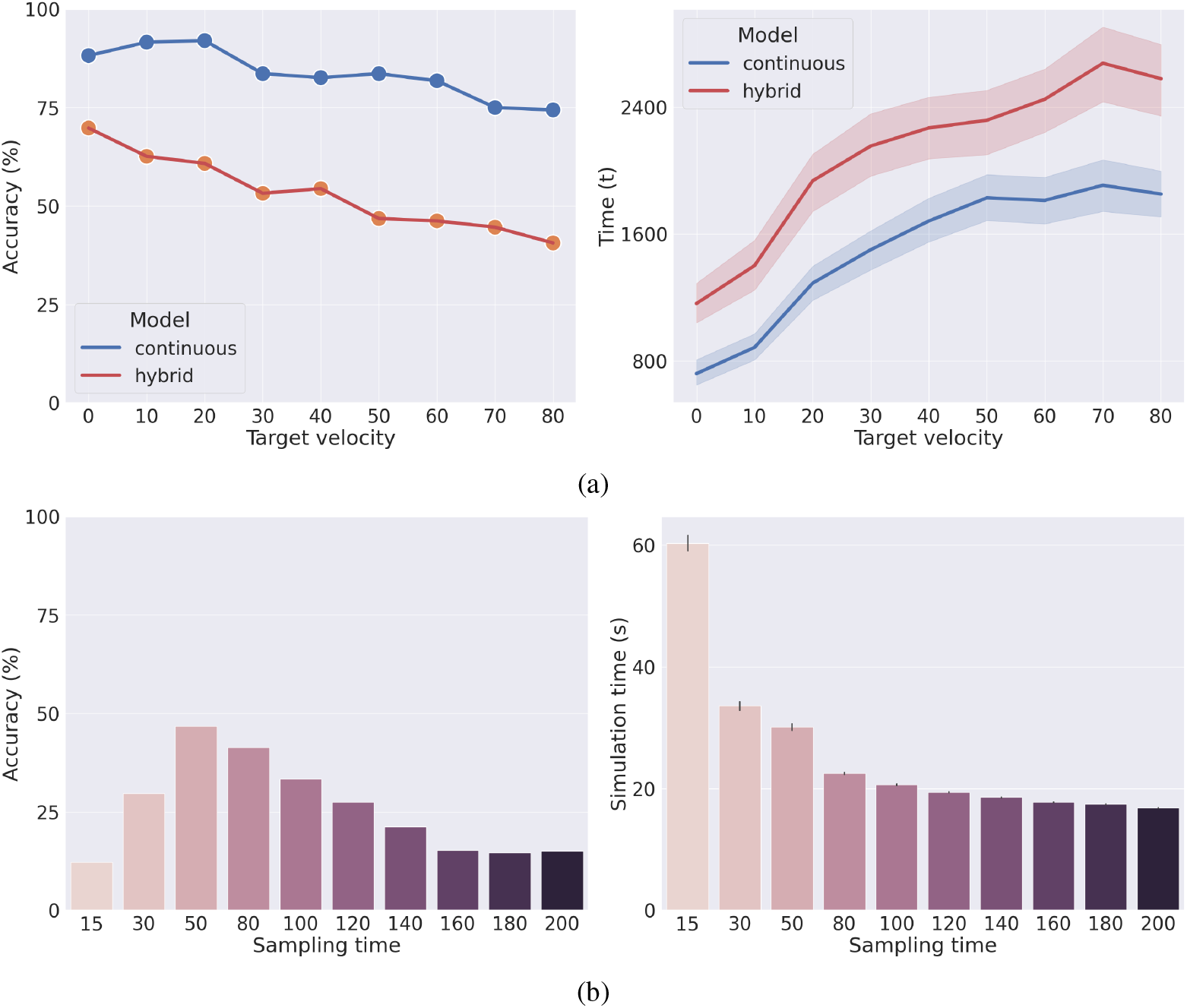
(a) Performance comparison between the hybrid and continuous-only models for different object velocities (left), and time needed to complete the task (right). Accuracy measures the number of successful trials. The right plot also shows the 95% confidence interval of every trial. As the object’s velocity increases, the grasping action becomes more difficult for the hybrid model since faster response planning is needed. Instead, the continuous-only agent adjusts its hidden causes based on the evidence accumulated at each time step. Despite using an a-priori (assumed to be learned) sequence of actions that does not permit replanning, this semi-flexible dynamics handles dynamic tasks more smoothly and efficiently. (b) Behavioral effects of the sampling time in the hybrid model with an object moving at a moderate velocity of 50 px/t. Accuracy (left) and average simulation time (right) are shown. Every condition is averaged over 500 trials. The simulation time is measured in seconds. The right plot also displays the 95% confidence interval of the time needed to complete the task.

In static grasping (i.e., zero object velocity), the continuous-only model achieves better performances compared to the hybrid model, with a difference in accuracy of 18.4%. This difference increases considerably when it comes to grasping a moving object, up to 33.8% for the highest target velocity. A similar difference can also be seen when considering the time needed to complete the task (the right plot of Figure 5a). We investigated the performance drop of the hybrid model in a second experiment illustrated in Figure 5b, showing a tradeoff between response time and computational complexity. In this experiment, we assessed the performances of the hybrid model by varying the sampling time window *T* from 15 to 200 time steps, with 500 trials per condition (i.e., 5000 trials in total). Here, we analyzed the agent’s accuracy and total simulation time, measured in seconds.

Greater sampling time allows the agent to accumulate more information and accurately estimate the discrete hidden states. However, longer sampling time also implies chunking the action into a smaller number of longer action primitives, which ultimately makes the grasping action fail since the object continuously changes its location during the sampling period. On the contrary, reducing the sampling time has the advantage of speeding up the agent’s planning process (i.e., faster reaction to environmental changes) so that it can grasp the object with greater accuracy, but only up to a certain point – after which there is a considerable performance drop. In this case, the agent would require longer policies since a reaching movement planned by the discrete model breaks down to a greater number of continuous trajectories. At the same time, since the discrete model is activated at a higher rate for action replanning, the simulation time steadily increases too, which can be associated with higher energy demands.

Figures 6a and 6b show the interplay between reaching and grasping actions in the hybrid and continuous models, using a constant sampling time window of 50 steps, and target velocity of 30 px/t. For both models, we initialized the target position and direction with the same values, as well as for the goal position. Here, five different phases can be distinguished: a pure reaching movement, an intermediate phase when the agent slowly approaches the object and prepares the grasping action, a grasping phase, another reaching movement and, finally, the object release. The computation of the action probabilities by the discrete model thus realizes a smooth transition between reaching and grasping actions, although at this stage still encoded as separate representations.

**Figure 6:**
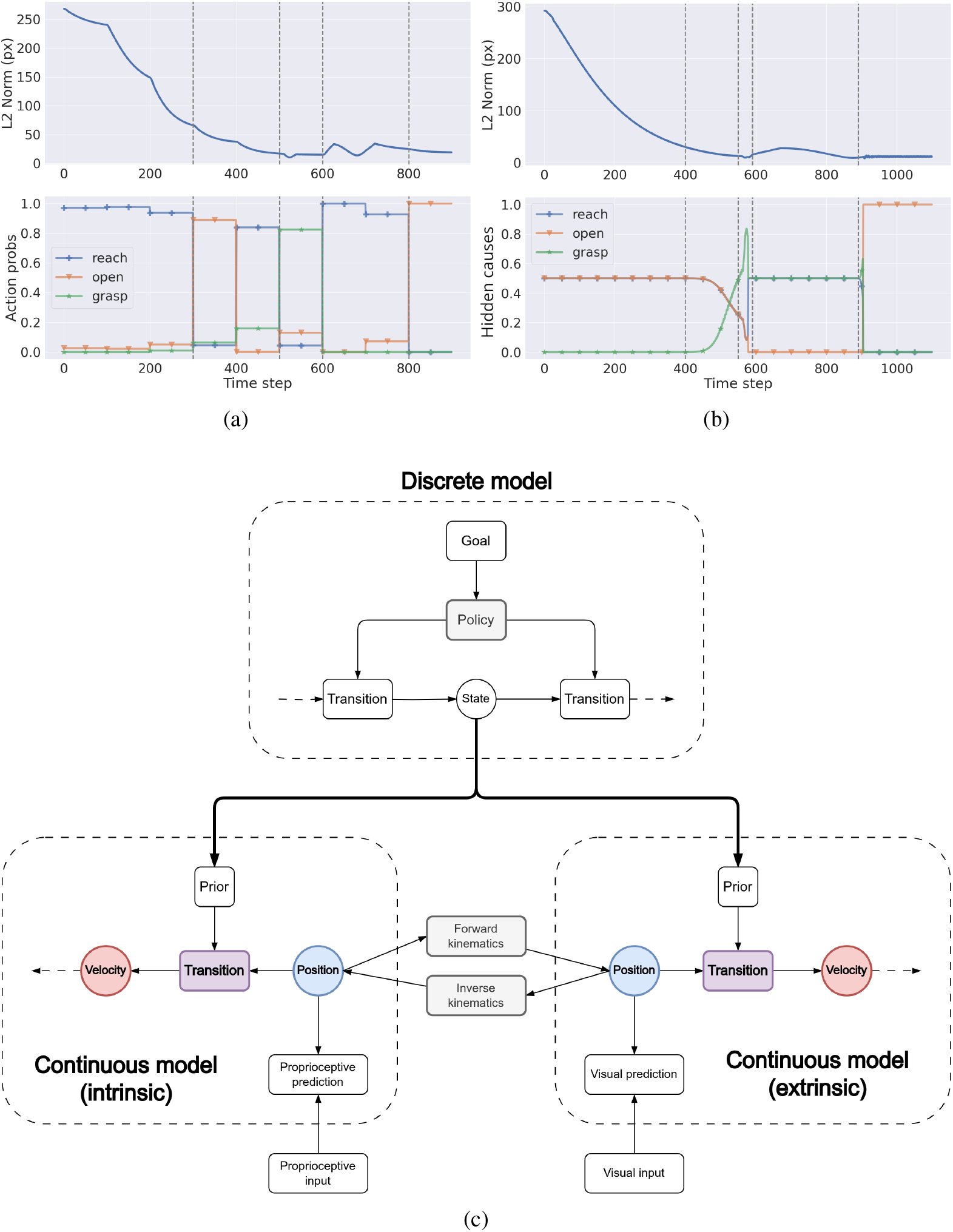
(a) Distance between hand and target (top) and action probabilities (bottom) for the hybrid model. (b) Distance between hand and target (top) and hidden causes (bottom) for the continuous model. (c) Graphical representation of the second associative phase. Continuous dynamics are represented in generalized coordinates of motion including position, velocity, acceleration, etc., allowing one to express with increasing fidelity the true environmental dynamics. Further, the outcomes of the discrete model not only bias the 0th temporal order (i.e., position) state, but steer the whole continuous trajectory toward a particular direction. In our reach-to-grasp task, this might correspond to considering the elementary actions of reaching and grasping as single discrete states that generate composite trajectories in the continuous domain.

This finding might provide a clue on how discrete actions lead to the emergence of smoother composite trajectories in the continuous domain, typical of intermediate phases of motor skill learning. In the second (associative) phase, the cortical activity begins to shift toward posterior areas. The process can be modeled as in Figure 6c, with a balanced work between discrete and continuous models in which the latter start to construct their own task dynamics, eventually leading to the model of Figure 3b. Note the differences with Figure 2, wherein the continuous models only receive a static prior resulting from a combination of all discrete states. In summary, while in the cognitive phase the task would be comprised of a reaching movement, an effort to find the next optimal action, and then a pure grasping action, in the associative phase the continuous models would handle the transition generating a fluid movement that makes the hand close as soon as it approaches the object. The discrete model could hence represent the task with a reaching-grasping action and a reaching-releasing action, with reduced computational demand.

## 4 Discussion

Understanding the mechanisms that support task specialization and motor skill learning is compelling for making advances with robots and intelligent systems. A major issue is that high-level and low-level processes are often analyzed and developed with different perspectives and techniques, as in optimal control algorithms [61] or deep neural networks [62]. Instead, the flexibility and robustness behind human and animal motor systems reside in that the learning of a new skill is a unified process that initially involves prevalent activation of prefrontal brain areas, and gradually shifts toward posterior and subcortical regions [12]. Indeed, constantly relying on the whole brain hierarchy is computationally demanding, and high-level areas are unable to track environmental changes rapidly. Highly dynamic tasks can be solved more efficiently by offloading the learned action transitions and policies to lower, faster hierarchical levels operating in continuous domains [63, 12]. For example, evidence suggests that repetitive and rhythmic movements do not involve activation of prefrontal areas but only rely on sensorimotor circuits [64]. Among those, the Posterior Parietal Cortex (PPC) is known for its involvement in reaching and grasping movements [65, 66] and encoding multiple goals in parallel during sequences of actions as in multi-step reaching [67], even when there is a considerable delay between goal states [68].

In this study, we show how the computations performed by hybrid and continuous models in active inference can be compared to different phases of motor skill learning. When the agent is required to interact with a novel situation, high-level planning is essential because the low levels alone cannot minimize the generated prediction errors, which then climb up the cortical hierarchy. As the agent practices the task and learns transitions that account for environmental uncertainty and possible dynamic elements, the prediction errors arising during the unfolding of the task can be explained away more efficiently by lower-level specialized neurons; as a result, repeated calling of high levels becomes redundant and ceases. Even using elementary discrete actions, a composite movement corresponding to an approaching phase between reaching and grasping naturally arises from Bayesian model reduction. This is due to the continuous evidence accumulation providing a smooth transition between discrete hidden states, as shown in Figures 6a and 6b. Therefore, we propose that the dynamics of a continuous model may have a specular structure to their discrete counterpart, that is, the final trajectory is generated by weighting independent distributions related to some action primitives. As a result, the composite movement might be embedded into the continuous dynamics with two parallel processes: while a discrete model can rapidly impose and infer specific trajectories, the dynamics precisions are additionally adapted through the same mechanisms of free energy minimization, so that the agent can score how well the action primitives perform. Importantly, the dynamics precisions act not just as modulatory signals but represent the agent’s confidence about the current state of the task – in a specular manner to sensory precisions [57]. The higher the precision of a specific trajectory, the more useful it is in explaining the agent’s final goal; on the other hand, a low trajectory precision means that the agent is not confident about it for a particular context. Nonetheless, this process does not mean that action primitives cannot be used anymore, but only that task-specific structures are constructed at low levels for the particular task considered. In fact, action primitives are maintained and can be used to create new combined trajectories to be applied to similar tasks.

Further, the results in Figure 5b show an interesting tradeoff between reaction times, planning capabilities, and computational demand. If the environment changes with high frequency, occasional calling of the discrete model does not permit to respond in time to novel sensory observations. On the other hand, periodic action replanning is counterproductive beyond a certain limit since, although allowing the agent to react more rapidly to environmental changes, it increases the effort of the discrete model to find the correct policy, as a higher number of discrete steps are needed. Instead, a continuous-only model achieves the best performances in less time and with minimal resource expenditure. However, this comes at the cost of being unable to adapt to changes in the hidden causes of reflexive control as inference when the context changes, resulting in an inability to plan in a context-sensitive manner.

The presented work introduces two novelties compared to state-of-the-art hybrid models. First, the reduced priors are generated dynamically at the beginning of each discrete step through the specification of flexible intentions – which roughly correspond to the discrete actions – enabling the agent to operate in dynamic environments [57]. Second, the discrete model can impose priors and accumulate evidence over multiple continuous modalities, e.g., intrinsic and extrinsic. This has the advantage that one can realize more complex goals (e.g., reaching with specific affordances, as shown in Figure 4b) and has access to more information for inferring the discrete hidden states (e.g., extrinsic for position, and intrinsic for hand status).

Finally, the analogies we presented between discrete, mixed, and continuous models may shed light on the information processing within cortical regions. It has been hypothesized that the cortex works within a system of discrete models of increasing hierarchical depth (e.g., motor, associative, limbic) [11], and various studies that recorded neural populations during generation of smooth movements seem to point at this direction [69, 70, 71]. Instead, the interface with the continuous models that receive sensory observations is supposed to be achieved through subcortical structures, such as the superior colliculus [35] or the thalamus [34]. Since deeper cortical levels are found to encode increasingly discretized representations [18], the latter could also be the consequence of the invariability of neural activity that results during inference when the hierarchy is spatially deep. Cortical computations have been simulated by continuous-time active inference [72, 73] too, but as well noted in [18], even the first computations of neural processing could be viewed as dealing with lots of little categories. In this view, whether it makes more sense to consider a specific level as discrete or continuous, from a high-level perspective a gradual transition between the two modalities could occur. Therefore, since all regions eventually have to deal with continuous signals, looking for analogies between the two processes might be helpful (e.g., the policy optimization of discrete models is generally compared to corticostriatal pathways [34], but reward-related activity has also been recorded in the Primary Visual Cortex V1 [74]).

Overall, our results provide some hints into how task specialization may occur in biologically plausible ways. However, in this preliminary study, the different phases were analyzed separately to show the relationships between policies and dynamic trajectories. Further studies are needed to understand how the generation of prediction errors throughout the hierarchy leads to the adaptation from discrete to continuous processes. Two promising directions would be to implement deep hierarchical models [55], and to hidden causes and dynamics precisions of the continuous models – which in our simulations were kept fixed throughout the task – adapt to the prediction errors, thus allowing self-modeling based on the current task and environmental state. Finally, future works will assess the capabilities of the hybrid active inference model to simulate human movements. For instance, a more realistic model would include a belief of real object dynamics, along with the dynamics desired by the agent. This would possibly account for anticipatory effects, which are required when modeling experiments that consist of catching moving objects on the fly, such as when baseball players try to intercept a thrown ball [75]. The proposed model may then be assessed against kinematic and neural data collected from humans and primates.

## Data availability

Code and data have been deposited in GitHub (https://github.com/priorelli/discrete-continuous).

## Acknowledgments

This research received funding from the European Union’s Horizon H2020-EIC-FETPROACT-2019 Programme for Research and Innovation under Grant Agreement 951910 “MAIA” to I.S. and the Italian National Recovery and Resilience Plan (NRRP) Project PE0000006, CUP J33C22002970002 “MNESYS” and Project PE0000013, CUP B53C22003630006, “FAIR”. The funders had no role in study design, data collection and analysis, decision to publish, or preparation of the manuscript.

